# A red-eye mutant in *Nesidiocoris tenuis* (Hemiptera: Miridae) is caused by exon skipping due to an indel mutation in the *scarlet* gene

**DOI:** 10.64898/2026.07.31.741938

**Authors:** Tomofumi Shibata, Kaoru Saeki, Chiharu Saito, Takuya Uehara

## Abstract

*Nesidiocoris tenuis* is an important zoophytophagous mirid bug used as a biological control agent in agriculture, and breeding efforts based on genomic information aim to increase its utility. Visible eye-color mutants are useful genetic markers because they are easily distinguishable and are therefore widely used in insect genetics and genome editing studies. Here, we investigated the genetic basis of a spontaneous red-eye mutant identified in a laboratory strain of *N. tenuis*. Classical crossing experiments suggested that the red-eye phenotype is controlled by a single recessive locus. RNA-seq and RNA interference (RNAi) analyses identified *scarlet* and *cinnabar* as the primary candidate genes associated with the phenotype. Further genomic analysis revealed a large deletion and insertion within exon 5 of the mutant *scarlet* allele, potentially causing exon skipping and disrupting transporter structure. The insertion pattern is consistent with a microhomology-mediated break-induced replication (MMBIR)/fork stalling and template switching (FoSTeS)-like event that may have been generated through polymerase *θ*-mediated repair. Together, these findings identify the causative mutation underlying the red-eye phenotype and provide a useful visible marker for future functional genetic studies and genome-assisted breeding in *N. tenuis*.

## 1. Introduction

*Nesidiocoris tenuis*, commonly known as the tomato bug, is an omnivorous species exhibiting both carnivorous and herbivorous feeding habits. This dual feeding strategy allows it to utilize both animal- and plant-derived food resources and makes it a promising biological control agent in agriculture worldwide (Pérez-Hedo and Urbaneja, 2016). *N. tenuis* preys on agricultural pests such as the silverleaf whitefly (*Bemisia tabaci*) (Sanchez, 2008) and the tomato leafminer (*Tuta absoluta*) (Shaltiel-Harpaz et al., 2016). However, depending on its population density, it may also damage crops through its herbivorous feeding habits (Calvo et al., 2009). Therefore, suppressing herbivorous feeding behavior is one of the primary goals of breeding programs for this species (Pérez-Hedo et al., 2024). More broadly, several undesirable traits limit the effectiveness of *N. tenuis* as a biological control agent, making genome-assisted breeding and functional genetic approaches increasingly important.

Recently, we generated a chromosome-level genome assembly of *N. tenuis* to provide a foundation for investigating the genetic basis of important traits such as its feeding habits (Shibata et al., 2024). However, genomic resources alone are insufficient to advance genome-assisted breeding. Additional foundational techniques, including genome editing and controlled inbreeding, are also needed. Easily distinguishable visible markers, such as eye-color mutants, may therefore provide valuable tools for applied genetic studies. Indeed, several studies developing genome-editing approaches in insects have targeted eye-color genes because the resulting mutant phenotypes are easy to recognize (Shirai et al., 2022; Reding et al., 2023).

During long-term maintenance of insect colonies under laboratory conditions, spontaneous mutants occasionally emerge. Eye-color mutations are among the most frequently reported mutations because they are easily distinguishable. Red-eye mutants have been reported in several hemipteran species, and the causative genes or genomic regions underlying these phenotypes have been identified. (Shimizu and Kawasaki, 2001; Snodgrass, 2002; Moraes et al., 2005; Liu et al., 2014; Reche et al., 2025). In our laboratory, we identified a spontaneous red-eye mutant in a population of *N. tenuis* originating in Shizuoka Prefecture, Japan **(Fig. 1)**, and subsequently established a stable red-eye mutant strain. Identifying the causative genes or genomic regions underlying these phenotypes. Determining the causative gene or genomic region responsible for the red-eye phenotype is essential for incorporating this strain into our breeding program.

**Figure 1.**
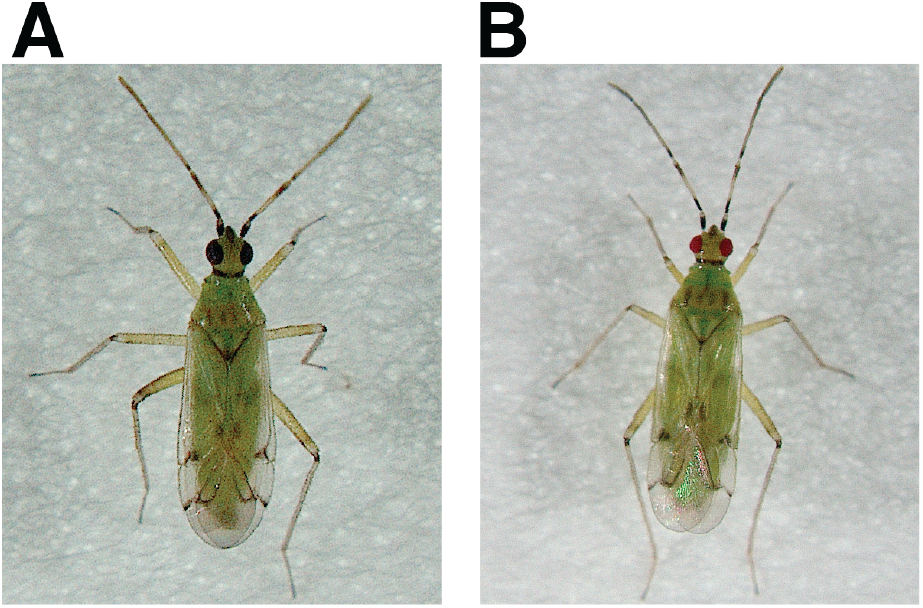
Wild-type and red-eye mutant *Nesidiocoris. tenuis*. **(A)**Adult male of wild-type *N. tenuis* originating in Shizuoka Prefecture, Japan. **(B)**Adult male of the red-eye mutant strain of *Nesidiocoris. tenuis*.

Here, we combined classical genetics, RNA-seq, RNA interference (RNAi), and genomic analyses to identify the causative mutation and characterize the insertion or deletion underlying our spontaneous red-eye mutant in *N. tenuis*. In addition, we discuss its potential applications for functional genetic studies and genome-assisted breeding.

## 2. Materials and Methods

### 2.1 Insects

Insects were reared on *Ephestia kuehniella* Zeller eggs (Ga-Ran; Agrisect Inc., Ibaraki, Japan) provided as a food source; the eggs were affixed to 18 × 50 mm adhesive labels. Several stems of *Plectranthus amboinicus* (Lamiaceae), trimmed to approximately 8 cm in length, were provided as oviposition substrates and a source of moisture. Insects were maintained at 25 ± 1°C, 50–70% relative humidity; under a 16:8–h light:dark photoperiod.

The Shizuoka strain of *N. tenuis* was established by combining individuals collected in Iwata City in 2020 (34.728°N, 137.826°E) and Kakegawa City in 2021 (34.682°N, 138.043°E) in Shizuoka Prefecture, Japan. The strain was maintained in a greenhouse on insectary plants established for conservation biological control. We identified a red-eye mutant in the Shizuoka strain and subsequently established a red-eye mutant strain. Briefly, unmated red-eye mutant males from the Shizuoka strain were crossed with red-eye mutant females from the same strain. Each pair was placed in a plastic case (138 × 109 × 53 mm), and sibling crosses were continued for more than two generations.

### 2.2 Genetic crosses

To determine the mode of inheritance of the red-eye phenotype, reciprocal crosses were performed between the red-eye mutant and wild-type strain. In the F0 generation, four pairs of virgin adult red-eye mutant males and wild-type females, and four pairs of virgin adult wildtype males and red-eye mutant females, were crossed to generate F1 progeny. In the F1 generation, eight mating pairs among progeny derived from crosses involving red-eye mutant males and 12 mating pairs of progeny derived from crosses involving red-eye mutant females were established to generate F2 progeny. Segregation ratios in the F2 generation were analyzed using a chi-square (*χ*^2^) test to determine the mode of inheritance according to Mendelian expectations. All genetic crosses were conducted under the rearing conditions described above.

### 2.3 RNA extraction and sequencing

Two male and two female *N. tenuis* individuals from each of the red-eye mutant and the wild-type strain, both derived from the Shizuoka strain, were used for total RNA extraction. Total RNA was extracted using an RNeasy Plus Mini Kit (Qiagen, Hilden, Germany). Approximately 50 ng/µL of RNA was obtained from each individual, as ameasured using a NanoDrop microvolume spectrophotometer (Thermo Fisher Scientific, MA, USA). Paired-end sequencing was performed on a NovaSeq 6000 system using a TruSeq Stranded mRNA Library Prep Kit (Illumina Inc., San Diego, CA, USA).

### 2.4 RNA-seq analysis

The quality of the raw-sequencing reads was assessed using FastQC v0.12.1 (https://www.bioinformatics.babraham.ac.uk/projects/fastqc/). Low-quality reads were filtered out using fastp v0.24.0 (Chen et al., 2018) with default parameters. The quality of the trimmed reads was re-evaluated using FastQC, confirming improved read quality. The processed reads were used for subsequent analyses. RNA-seq reads were mapped to the Japanese reference strain genome (Shibata et al., 2024) using STAR v2.7.11b (Dobin et al., 2013), with a unique mapping rate of approximately 80% for each sample. Gene-level expression values were quantified using RSEM v1.3.3 (Li and Dewey, 2011), and isoform-level expression data were used for differential gene expression analysis, which was conducted using DESeq2 v1.46.0 (Love et al., 2014) in R v4.2.0 with RStudio v1.1.456.

Assignment of *N. tenuis* ommochrome pathway genes was based on comparison with the eye-color genes *cinnabar, scarlet, cardinal, white, karmoisin*, and *vermilion*, from *Lygus hesperus* Knight (Brent and Hull, 2019), using a BLASTX top-hit search. Volcano plots were generated using EnhancedVolcano v1.24.0 (Blighe et al., 2019). Expression levels of six candidate ommochrome pathway genes were evaluated using log_2_-transformed transcripts-per-million (TPM) values (log_2_[TPM + 1]) and compared between the wild-type and red-eye mutant strains using a twosided Mann-Whitney *U* test. Correction for multiple testing across the six genes was performed using the Benjamini-Hochberg false discovery rate (FDR) procedure.

### 2.5 Gene silencing assay using RNA interference

We designed 200 -500 bp primers for RNAi (**Supplementary Table S1**). Double-stranded RNA (dsRNA) was synthesized using a Promega T7 Ribo-MAX Express RNAi System Kit (P1700; Promenga, Madison, WI, USA), and the dsRNA concentration was diluted to 500 ng/µL. An aliquot (1 µL) of dsRNA was injected into the thorax or abdomen of third-instar nymphs. After eclosion, adult eye color was examined and target gene expression levels were quantified by qPCR using Thunderbird Next SYBR qPCR Mix (QPX-201; Toyobo Co. Ltd., Osaka, Japan). Statistical significance was evaluated using Fisher’s exact test against the EGFP control. *P*-values were adjusted for multiple comparisons using the Benjamini-Hochberg FDR correction.

### 2.6 Genome analysis and protein reconstruction

After narrowing down the candidate genes responsible for the red-eye mutation, genome-mapped RNA-seq data were manually inspected using IGV v2.19.1 (Thorvaldsdóttir et al., 2013). Primers were designed to amplify a 921 bp region encompassing the target site for Sanger sequencing (**Supplementary Table S1**). Cycle sequencing was performed using BigDye Terminator v1.1 (Thermo Fisher Scientific). Raw sequencing data were base-called using Sequencing Analysis Software v7.0 (Thermo Fisher Scientific), and low-quality bases were removed using Sequencher v5.0 (Gene Codes Corporation, Ann Arbor, MI, USA). Trimmed reads were aligned using MAFFT v7 (Katoh and Standley, 2013), and the mutation site was identified.

Potential exonic splicing enhancers (ESEs) were identified using ESEfinder 3.0 (Cartegni et al., 2003). The predicted structure of the protein encoded by the candidate gene was generated using ColabFold v1.5.3 (Mirdita et al., 2022). To identify sequence features associated with potential DNA double-strand break (DSB) formation, the full-length *scarlet* genomic region was obtained from the genome assembly of the Japanese reference strain. The sequence surrounding the predicted breakpoint was analyzed for local sequence composition. AT content was calculated using a custom Python script with a sliding window of 100 bp and a step size of 10 bp. Poly(dA:dT) tracts ( ≥15 consecutive A or T nucleotides) were identified within each window and plotted across the analyzed region.

## 3. Results

### 3.1 Genetic cross supports single-gene Mendelian inheritance

To investigate the inheritance pattern of the red-eye phenotype, pairs consisting of black-eye and red-eye individuals were established as the F0 generation. All 70 F1 individuals obtained from these crosses had the black-eye phenotype, indicating that the red-eye mutation is recessive. To produce the F2 generation, eight mating pairs derived from the F1 progeny of crosses involving red-eye males, and 12 mating pairs derived from the F1 progeny of crosses involving red-eye females, were established. Among the 190 F2 individuals obtained, 51 had the red-eye phenotype and 139 had the black-eye phenotype.

To evaluate these results, we tested three models of inheritance of the red-eye mutation; a single recessive gene, complementary genes, and two recessive genes **(Table 1)**. The null hypotheses for the complementary gene and two recessive gene models were rejected (*P* < 0.001, *χ*^2^ test), whereas that for the sigle recessive gene model was not (*P* = 0.34, *χ*^2^ test), indicating that the red-eye phenotype is most likely controlled by a mutation in a single gene.

**Table 1.** Chi-square test of segregation ratios under different genetic models. Observed wild-type mutant ratio in the red-eye F2 generation was 139:51.

| Cross | Genetic hypothesis | Expected ratio | Observed ratio | $\chi^2$ | df | *P*-value |
| --- | --- | --- | --- | --- | --- | --- |
| F<sub>1</sub> | Single recessive gene | 1:0 | 70:0 | - | - | - |
| F<sub>2</sub> | Single recessive gene | 3:1 | 139:51 | 0.34 | 1 | 0.56 |
| F<sub>2</sub> | Complementary genes | 9:7 | 139:51 | 22.07 | 1 | 2.63e-06 < 0.001 |
| F<sub>2</sub> | Two recessive genes | 15:1 | 139:51 | 137.50 | 1 | 9.37e-32 < 0.001 |

### 3.2 Ommochrome pathway genes are associated with the red-eye mutant phenotype

Because red-eye phenotypes in insects are frequently associated with genes involved in the ommochrome pathway **(Fig. 2A)**, RNA-seq analysis was performed to compare gene expression profiles between the redeye mutant and the wild-type strains. Principal component analysis (PCA) showed that sex-related variation was more pronounced than strain-related variation, although transcriptomic differences between the strains were also evident **(Fig. 2B)**. Genome-wide differential expression analysis did not reveal a clear overall pattern of differential expression among the ommochrome pathway genes **(Supplementary Fig. S1)**. Therefore, the expression levels of individual genes involved in the ommochrome pathway were examined in detail. The expression levels of *cinnabar, scarlet, cardinal*, and *white* were reduced in the red-eye strain relative to the wild-type strain; however, after FDR correction, none of these genes remained significantly differentially expressed **(Fig. 2C)**.

**Figure 2.**
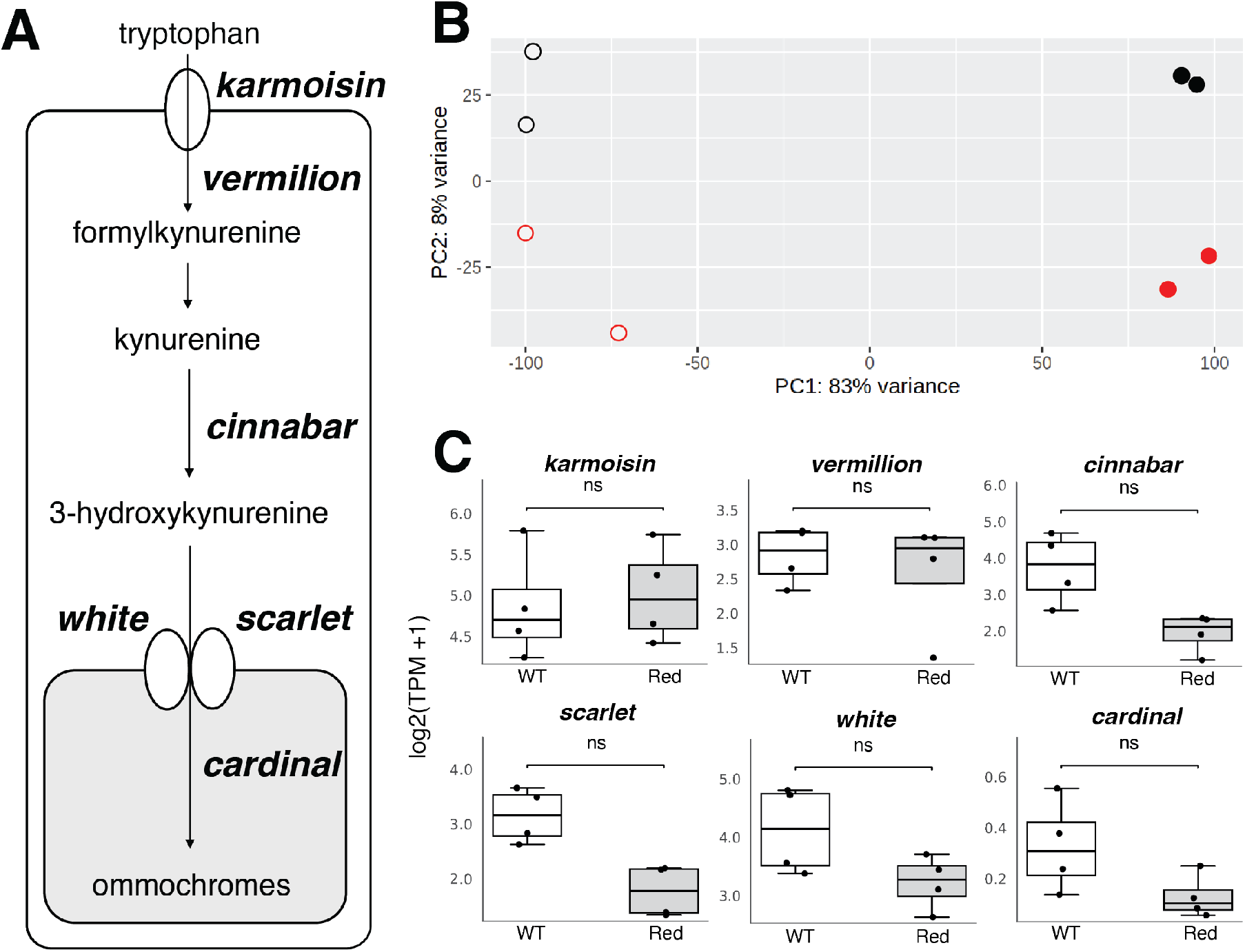
Overview of the ommochrome biosynthetic pathway and Differential Gene Expression analysis between the red-eye and wild-type strains using DESeq2. (**A**) Over view of the ommochrome biosynthetic pathway. Gene names are shown in ***bold italics***; chemical compounds are shown in regular font. (**B**) Principal component analysis of RNA-seq data. Black filled circles, wild-type males; black open circles, wild-type females; red filled circles, red-eye mutant males; and red open circles, red-eye mutant females. (**C**) Log_2_ -transformed TPM values (log_2_ [TPM + 1]) for six candidate ommochrome pathway genes. wt, wild type; red, red-eye mutants. Points represent biological replicates. Differences in expression were assessed using a two-sided Mann-Whitney U test followed by Benjamini-Hochberg false discovery rate correction. ns, not significant.

### *3*.*3 Cinnabar* or *scarlet* is associated with the redeye phenotype

Based on the differential gene expression analysis results, *cardinal, cinnabar, scarlet*, and *white* were identified as candidate genes for further analysis. However, because *white* is involved in both the ommochrome and pteridine pathways (Vargas-Lowman et al., 2019), knockdown of this gene is expected to result in a white-eye phenotype rather than a red-eye phenotype. Therefore, the remaining three ommochrome pathway genes– *cinnabar, scarlet*, and *cardinal*–were selected for knock-down by RNAi. Following dsRNA injection into nymphs, expression levels of all three target genes were reduced relative to control (EGFP) **(Fig. 3A–C)**. Representative images showing the eye-color phenotypes after RNAi are shown in **Fig. 3D, E**. Many individuals injected with dsRNA targeting *cinnabar* or *scarlet* dsRNA exhibited a red-eye phenotypes after molting to adulthood **(Fig. 3E)**. In contrast, individuals injected with *cardinal* dsRNA generally exhibited an eye-colortion similar to that of wild-type when observed with the naked eye, although slight red pigmentation was observed at the eye margins under a stereomicroscope **(Fig. 3E)**. No individuals with the target phenotype were observed in the EGFP control group (0/9). In contrast, a red-eye phenotype was observed in 7 of 9 individuals (77.8%) following *cinnabar* knockdown and in 5 of 6 individuals (83.3%) following *scarlet* knockdown. The frequency of the redeye phenotype was significantly higher in both knock-down groups than in the control group (FDR-adjusted *P* = 0.0034). Following *cardinal* knockdown, the red-eye phenotype was observed in 2 of 10 individuals (20.0%), but its frequency did not differ significantly from that in the control group (FDR-adjusted *P* = 0.4737).

**Figure 3.**
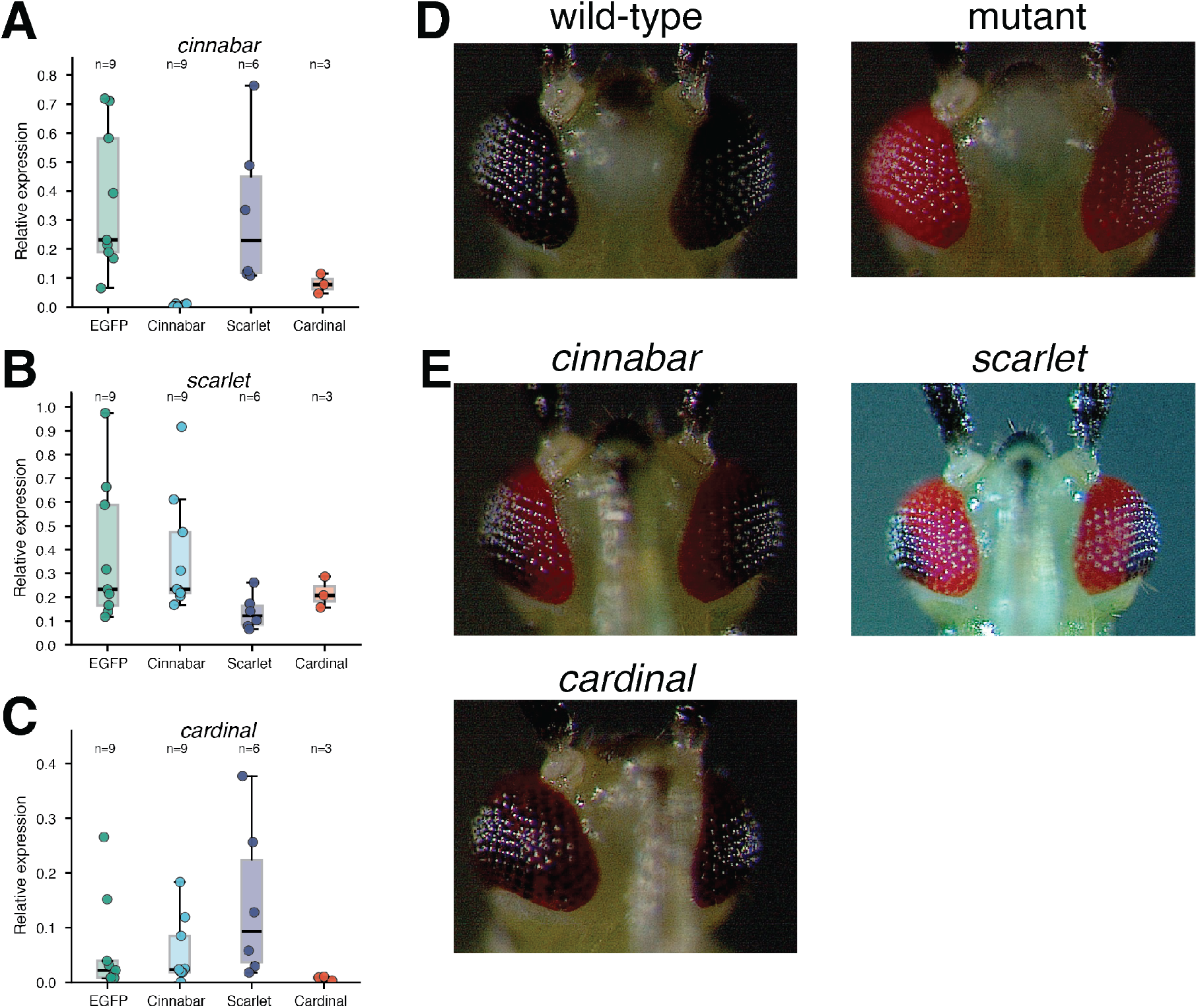
Relative expression levels of three ommochrome pathway three genes in control (EGFP) and RNAi-treated individuals, representative images of the resulting phenotypes. (A–C) Box plots showing target gene expression levels relative to *SOD*. Center lines indicate medians, boxes indicate interquartile ranges, and whiskers indicate the full data range. Sample sizes (*n*) are shown above each group. (A) *cinnabar* knockdown, (B) *scarlet* knockdown, and (C) *cardinal* knockdown. (D) Wild-type and red-eye mutant phenotypes.(E) Representative phenotypes following RNAi treatment targeting the indicated genes.

### 3.4 Mutation site in the *scarlet* gene of the red-eye mutant

Gene silencing of *cinnabar* or *scarlet* recapitulated the red-eye phenotype of the mutant *N. tenuis* strain. However, because knockdown of both genes produced a similar phenotypes, these results alone did not identify the causative gene. We therefore next examined the gene structures of these two candidate genes.

Mapping RNA-seq reads from the red-eye mutant and wild-type strains to the genomic regions containing these genes revealed reduced coverage across exon 5 of *scarlet* in the mutant strain **(Fig. 4A)**. In contrast, no clear differences in mapping patterns were observed for the other ommochrome pathway genes, suggesting that *scarlet* is the causative gene underlying the redeye phenotype. The predicted protein structure corresponding to the exon-skipped mRNA **(Supplementary Fig. S2)** showed fewer transmembrane *α*-helices, suggesting reduced or abolished transporter function **(Fig. 4B)**.

**Figure 4.**
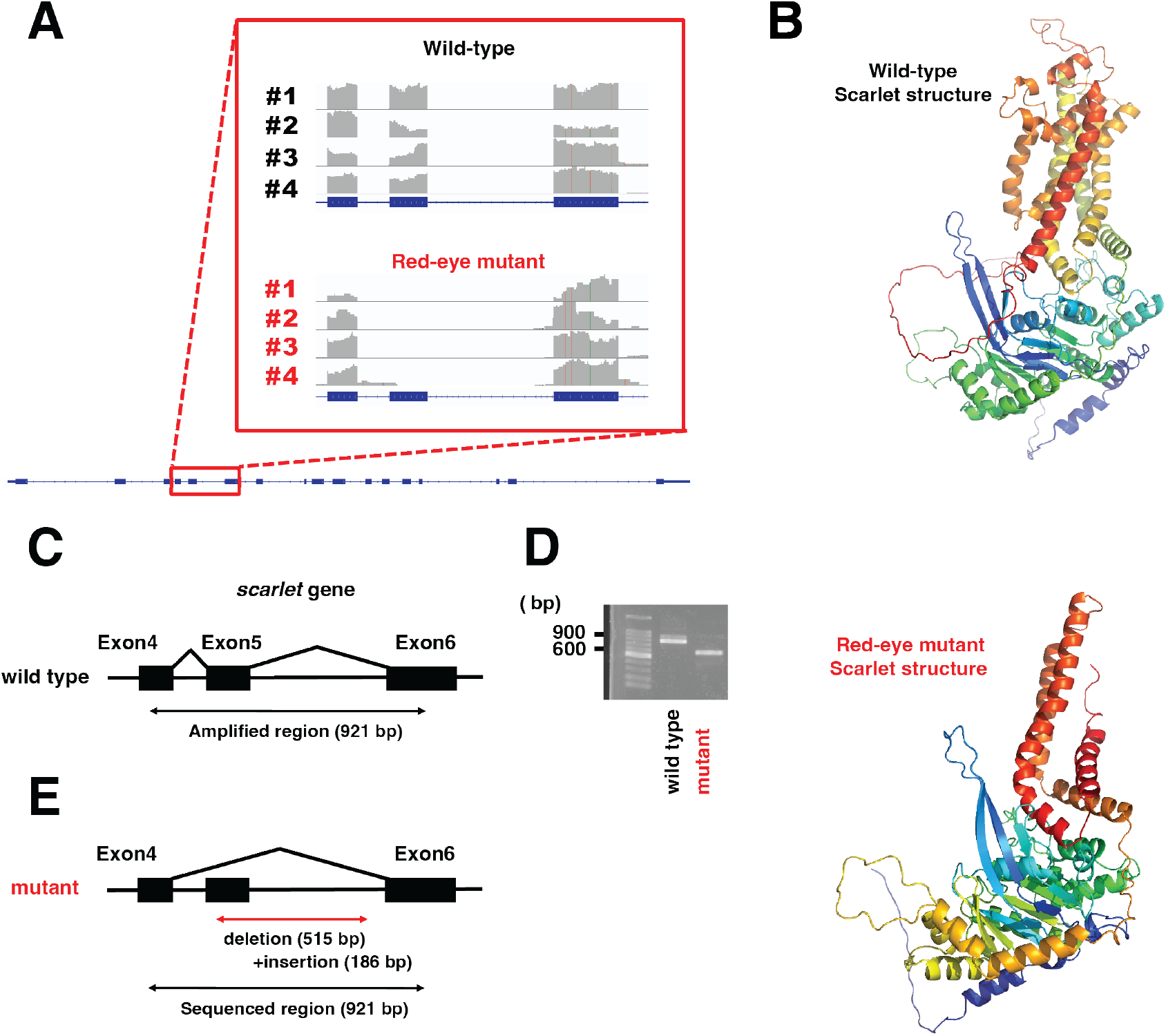
Gene expression pattern and indel mutation identified in the *scarlet* gene. (**A**) Visualization of mapped RNA-seq reads mapped to *scarlet* gene region. The region showing reduced coverage is highlighted by the red box. Labels #1–#4 indicate individual samples. (**B**) Predicted protein structures generated using ColabFold. Rainbow coloring represents the N-to C-terminal direction (from start codon to stop codon).The spiral-like structures in the upper right of the models are transmembrane *α*-helices. (**C**) Magnified view of the *scarlet* gene region highlighted by the red box in (**A**), spanning exons 4-6 in the wild-type strain. (**D**) PCR amplification of the region shown in (**C**), followed by agarose gel electrophoresis with a 500-bp ladder. (**E**) Schematic representation of the indel mutation in the mutant *scarlet* allele identified by Sanger sequencing.

PCR amplification of the 921-bp region of the *scarlet* gene encompassing exon 5 produced an approximately 900-bp fragment in wild-type individuals and an approximately 600-bp fragment in red-eye mutant individuals **(Fig. 4C, D)**. To characterize the indel mutation in the *scarlet* gene, Sanger sequencing was performed using the same amplified region. Sequence analysis revealed that the mutant *scarlet* allele contained a 515-bp deletion and a 186-bp insertion **(Fig. 4E, Supplementary Fig. S3)**. ESEs were predicted for the complete exon 5 sequence of the wild-type allele and the indel-containing exon 5 sequence of the mutant allele. The mutant allele contained fewer predicted ESEs than the wild-type allele, and the insertion region contained almost no predicted enhancers **(Supplementary Fig. S4)**.

## 4. Discussion

To identify the genetic basis of a spontaneous red-eye mutant in *N. tenuis*, we combined genetic, functional, and bioinformatic approaches. Our results demonstrated that the red-eye phenotype is controlled by a single autosomal recessive locus. Gene-silencing experiments narrowed the candidate genes to *cinnabar* and *scarlet*, while genomic and transcriptomic analyses identified *scarlet* as the causative gene. The mutant allele was found to contain a deletion encompassing the exon 5 of *scarlet*, and the sequence features surrounding the deletion suggest that it originated from a DNA damage followed by repair. Together, these findings establish *scarlet* as the gene responsible for the red-eye phenotype in *N. tenuis*.

Genetic analyses demonstrated that the red-eye phenotype is inherited as a single autosomal recessive trait, consistent with previous studies in Hemiptera (Shimizu and Kawasaki, 2001; Snodgrass, 2002; Liu et al., 2014). The red-eye strain established in this study has been maintained as a stable colony in the laboratory for several generations, suggesting that the mutation is not associated with major fitness costs under rearing conditions. This observation is consistent with findings in several insect species, where eye-color mutations generally have limited effects on viability under laboratory conditions (Snodgrass, 2002; Liu et al., 2014; Reche et al., 2025). However, it remains possible that the red-eye trait is disadvantageous under natural conditions. For example, the retina of a red-eye mutant in *Rhodnius prolixus* is susceptible to ultraviolet-induced damage (Insausti et al., 2013), suggesting that altered eye pigmentation may affect responses to short-wavelength light. This possibility is particularly relevant to the behavioral ecology of natural enemies. Our group has established techniques of manipulating the behavior of natural enemies through visual or olfactory signals (Ogino et al., 2016; Tokushima et al., 2016; Uehara et al., 2019a,b). In particular, short-wavelength light, including violet and ultra-violet light, has been shown to influence their behavior (Uehara et al., 2019a; Hall et al., 2021; Park and Lee, 2021). Therefore, an important direction for future research will be to determine whether the red-eye mutation influences visual sensitivity or behavioral responses to these light stimuli.

Based on differential gene expression analysis between the wild-type and red-eye mutant strains, we selected three ommochrome pathway genes –*cinnabar, scarlet* and *cardinal*– as targets for RNAi assays. Among these candidates, only *cardinal* knockdown produced a phenotype that largely resembled the wild-type **(Fig. 3D, E)**. Previous studies have suggested that ommochrome pigments can be produced through spontaneous redox reactions even in the absence of *cardinal* activity (Figon and Casas, 2019; Heu et al., 2022), which may explain the relatively weak phenotype observed following *cardinal* knockdown in the present study. Interestingly, knockdown of either *cinnabar* or *scarlet* was accompanied by reduced *cardinal* expression **(Fig. 3A, B)**. Although the mechanism underlying this association remains unknown, this observation may reflect transcriptional or metabolic interactions among genes in the ommochrome pathway. However, the regulatory relationships among these genes remain unclear and require further investigation.

The disrupted exon 5 region of *scarlet* exhibited markedly reduced RNA-seq coverage in the red-eye strain **(Fig. 4A)**. Subsequent Sanger sequencing revealed a complex structural variant consisting of a 515-bp deletion accompanied by a 186-bp insertion within exon 5 **(Fig. 5A)**. The 186 bp insertion showed no significant similarity to sequences in either public databases or the *N. tenuis* genome based on BLASTn searches, suggesting that it is unlikely to be derived from a known transposable element. Sequence comparisons suggested that the inserted sequence consists of three components: (i) a region showing limited similarity to nearby sequences (35 bp, 82% identity) **(Fig. 5B)**, (ii) an AT-rich low-complexity region with little detectable homology to adjacent sequences (82 bp, 89% AT content) **(Fig. 5C)**, and (iii) a region nearly identical to a flanking sequence and consistent with a microhomology-mediated break-induced replication (MMBIR)/fork stalling and template switching (FoSTeS)-like template-switching event (69 bp, 98% identity) **(Fig. 5D)**.

**Figure 5.**
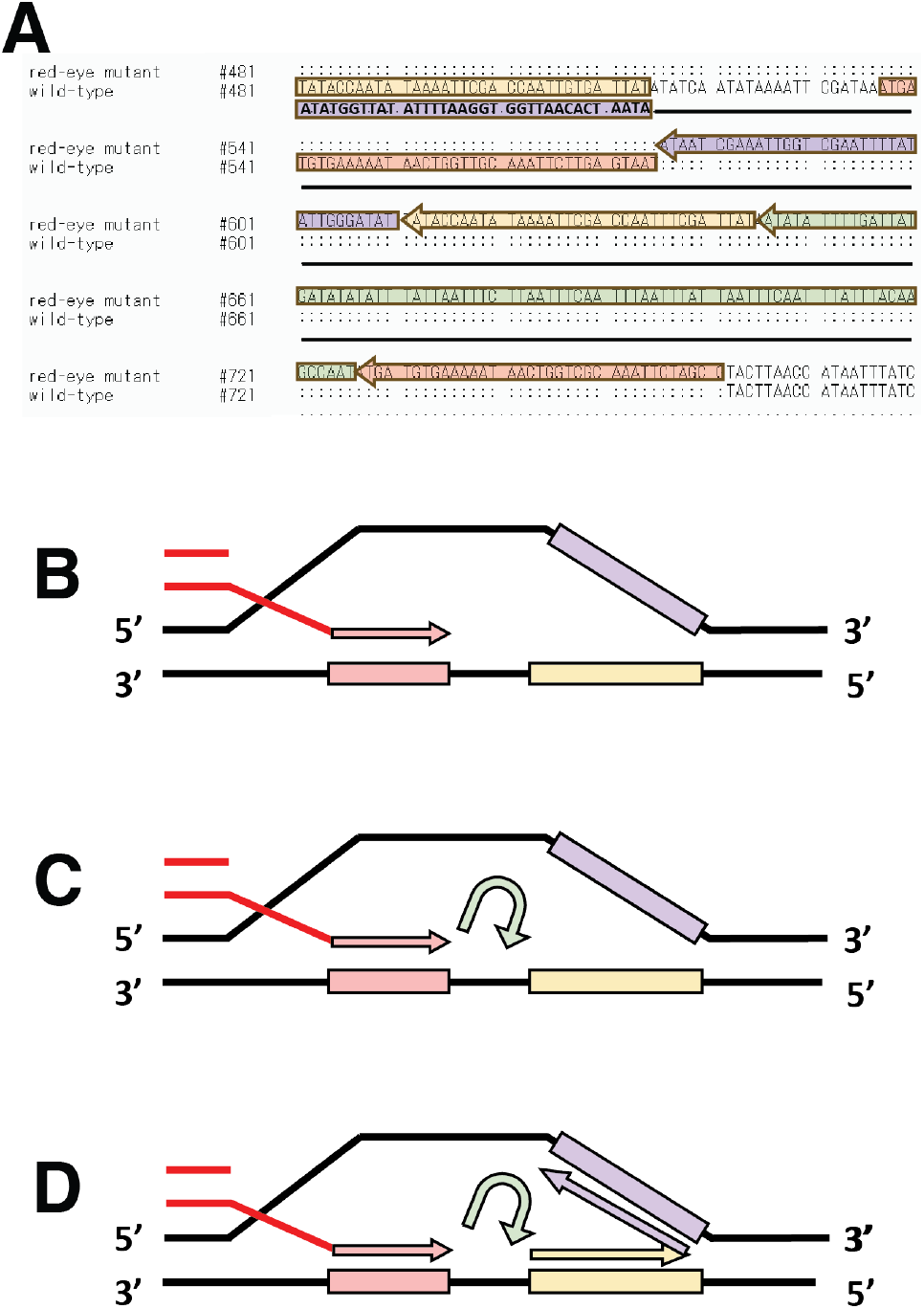
Proposed repair mechanism underlying the complex insertion in the mutant *scarlet* allele. **(A)**Schematic representation of the inserted sequence (excerpted from Supplementary Fig. S2). Arrows indicate the inferred direction of repair, and colors correspond to the regions highlighted by boxes of the same color. The region in the purple box is complementary to that in the yellow box. (**B**) Region potentially generated through repair and showing limited sequence similarity. (**C**) AT-rich, low-complexity region potentially generated by template-independent repair. (**D**) Region potentially generated through an MMBIR/FoSTeS-like template-switching event.

One possible explanation for the origin of this complex insertion is a template-switching event resembling MM-BIR (Hastings et al., 2009) or FoSTeS (Lee et al., 2007) during polymerase *θ*-mediated end joining (TMEJ), a DNA double-strand break repair pathway conserved across eukaryotes (Wood and Doublié, 2016; Marygold et al., 2020). Polymerase *θ* can repair DNA using microhomology and is known to generate complex insertion patterns through template switching or template-independent extension (Black et al., 2016; Yang and Gao, 2018). In particular, template-independent repair frequently produces low-complexity sequences, including AT-rich regions (Carvajal-Garcia et al., 2020). Therefore, the insertion structure observed here is consistent with repeated template-switching events during polymerase *θ*-mediated repair, resembling an MM-BIR/FoSTeS-like mechanism.

A prerequisite for this hypothesis is the occurrence of a DNA DNA double-strand break (DSB). AT-rich sequences and poly(dA:dT) tracts have been associated with genomic instability and increased susceptibility to DSB formation (Tubbs et al., 2018; Li and Wu, 2020). The indel identified in this study occurred within an AT-rich region of the *scarlet* gene containing multiple poly(dA:dT) tracts **(Supplementary Fig. S5)**. These sequence features are therefore consistent with a genomic region susceptible to DSB formation and may have facilitated the observed complex indel, although its exact origin remains unresolved. If this hypothesis is correct, our findings may represent one of the first exaples implicating an MMBIR/FoSTeS-like rearrangement as the causative mutation underlying a naturally occurring insect mutant. More importantly, the extensive disruption of exon 5 substantially reduced the number of predicted exonic splicing enhancers (ESEs) **(Supplementary Fig. S4)**. Together with the near absence of RNA-seq coverage across the affected exon, these findings strongly suggest aberraant splicing, potentially involving exon 5 skipping. Such splicing disruption would be expected to impair the normal structure and transporter function of the protein encoded by *scarlet* **(Fig. 4B)**.

Taken together, our results strongly support *scarlet* as the primary causative gene underlying the spontaneous red-eye phenotype in *N. tenuis*. Identification of the causative gene also expands the potential utility of this mutant strain in breeding and genetic studies. Genome-assisted breeding in insects remains at an early stage of development, and in *N. tenuis*, breeding efforts have largely relied on conventional selective approaches (Pérez-Hedo et al., 2024). Because the red-eye phenotype is easily distinguishable and the mutant strain can be stably maintained under laboratory conditions, this strain may provide a useful visible marker for future genome editing and functional genetic studies. Furthermore, if genomic variants tightly linked to agriculturally beneficial traits are identified near the *scarlet* locus, the red-eye phenotype could serve as an easily scored visible marker in marker-assisted breeding programs. However, as detailed behavioral and ecological evaluations have not yet been performed, future studies should determine whether this phenotype affects other fitness-related traits.

## Funding souces

This project was supported by JSPS KAKENHI (20H02992) and in part by the Cabinet Office, Government of Japan, Cross-ministerial Moonshot Agriculture, Forestry and Fisheries Research and Development Program (JPJ009237).

## CRediT authorship contribution statement

Tomofumi Shibata: Conceptualization, Validation, Formal analysis, Investigation, Writing - Original Draft, Writing - Review & Editing. Kaoru Saeki: Investigation, Writing - Review & Editing. Chiharu Saito: Resources, Writing - Review & Editing. Takuya Uehara: Conceptualization, Validation, Investigation, Writing - Original Draft, Writing - Review & Editing, Supervision, Funding acquisition.

## Acknowledgments

We thank Dr. T. Fukagawa of the National Research Institute of Police Science for valuable advice on RNA-seq analysis. We also acknowledge the efforts of the Insect Design Technology Group for maintaining the insect colony.

## Data availability

All raw reads obtained for RNA-seq analysis have been deposited in the DNA Data Bank of Japan under BioProjectID PRJDB42534 and Sequence Read Archive (SRA) under DRR1062891-DRR1062898.

## Supplementary Information

**Figure S1.**
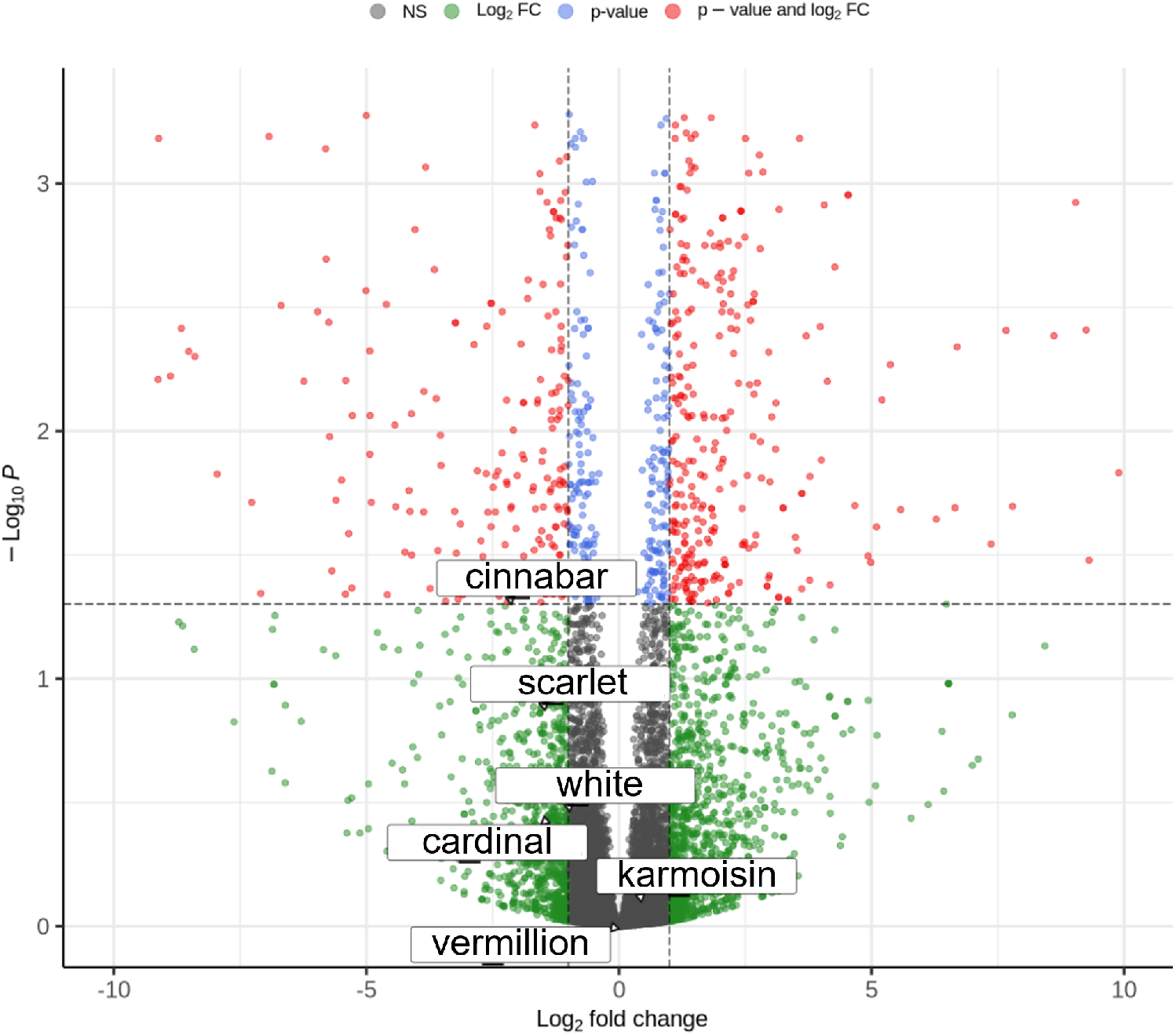
Volcano plot comparing the wild-type and red-eye mutant strains. *cinnabar* showed a significant adjusted *P* -value and exceeded the log2 fold change cutoff. *scarlet, white*, and *cardinal* exceeded the log2 fold-change cutoff but did not show statistically significant adjusted *p*-values. *karmoisin* and *vermilion* showed no significant differences in either adjusted *P* -values or log2 fold changes. The log2 fold-change cutoff was set at 1.0 and the adjusted *P* -value cutoff was set at 0.05.

**Figure S2.**
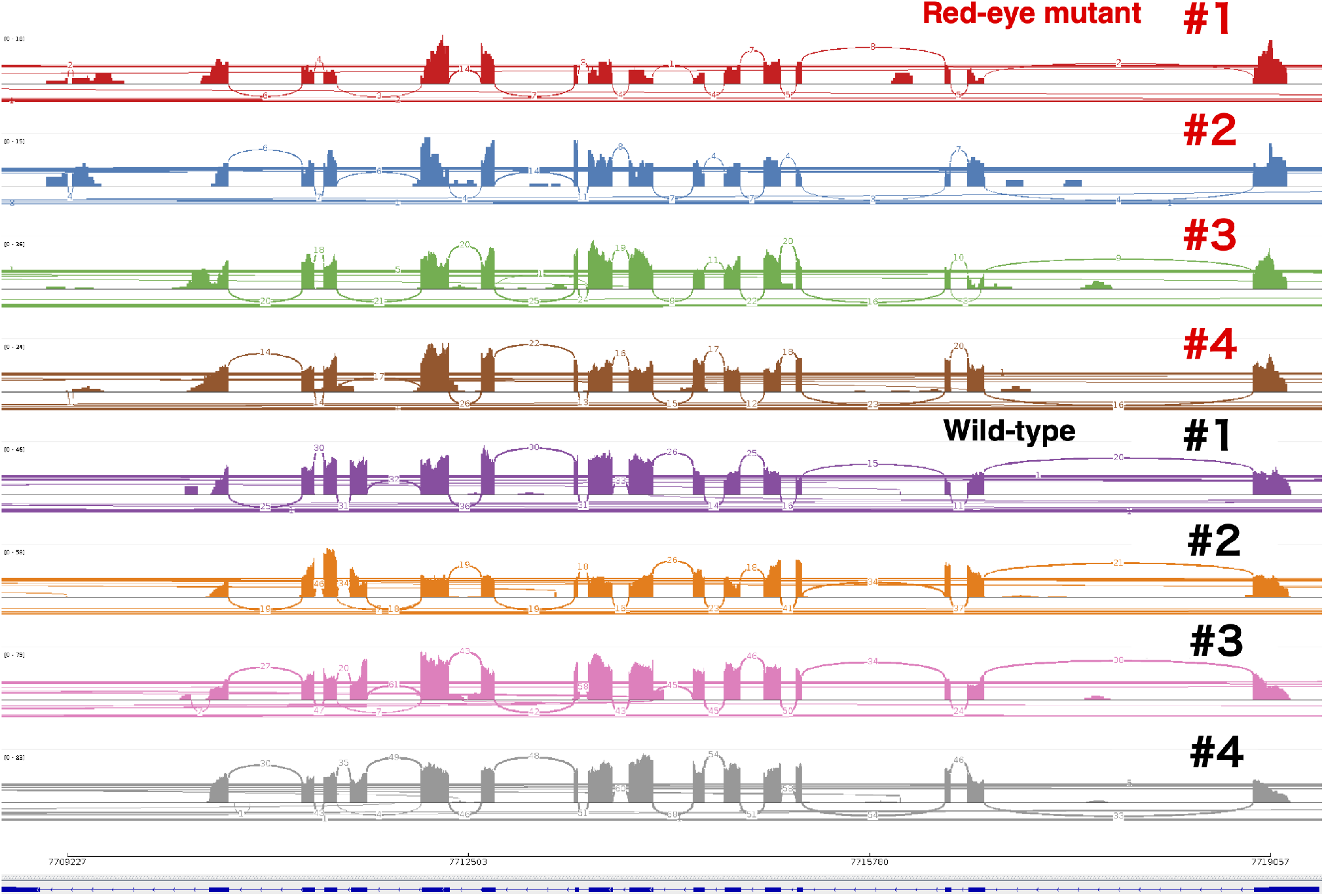
RNA-seq splice junctions across the *scarlet* gene shown as a sashimi plot. RNA-seq splice junctions across the entire *scarlet* gene are shown. Numbers on the arcs indicate the numbers of junction-spanning reads supporting each splice junction.

**Figure S3.**
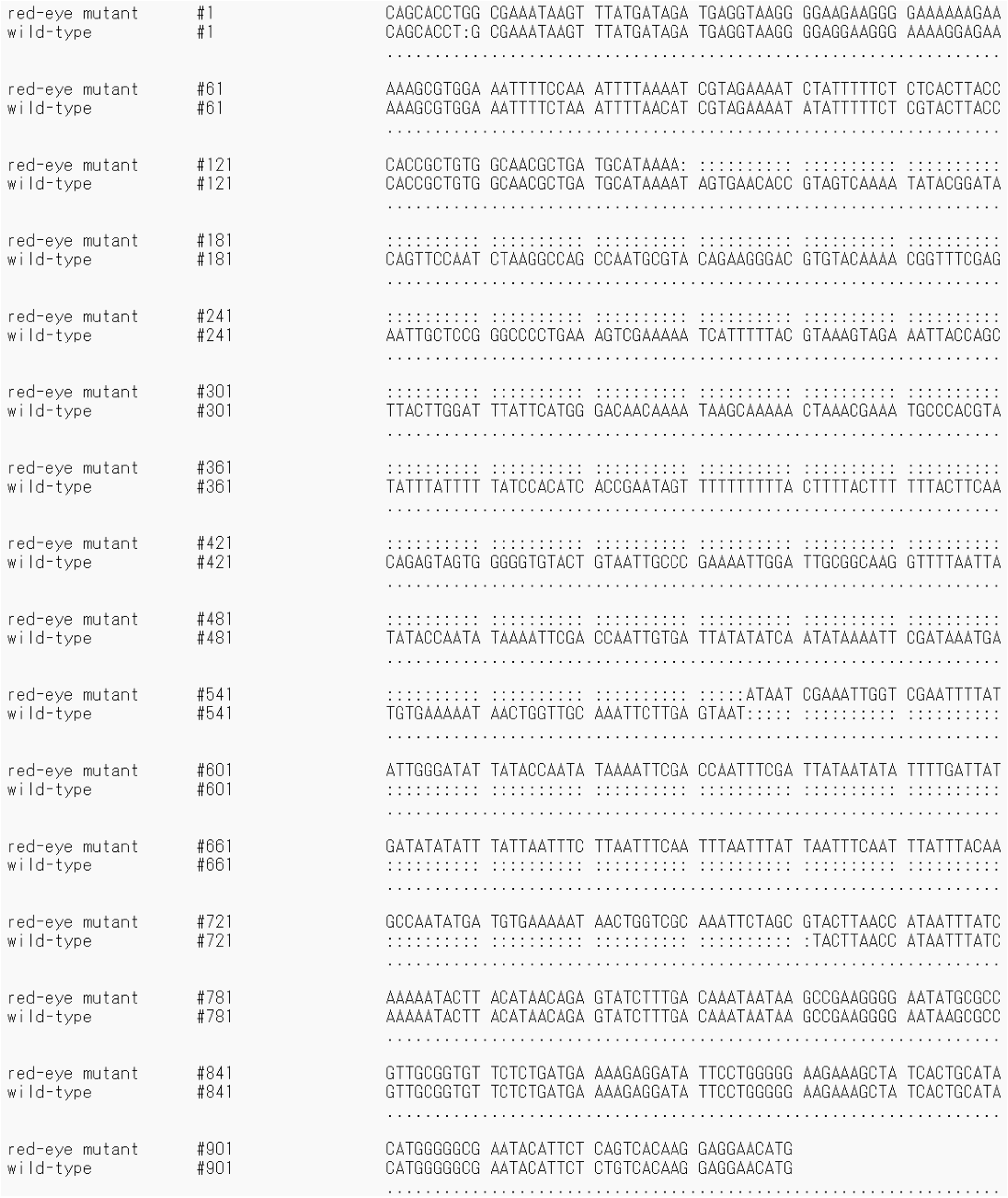
Comparison of Sanger sequencing data between the red-eye mutant and wild-type individuals. Colons indicate a region that could not be aligned between the red-eye mutant and wild-type sequences.

**Figure S4.**
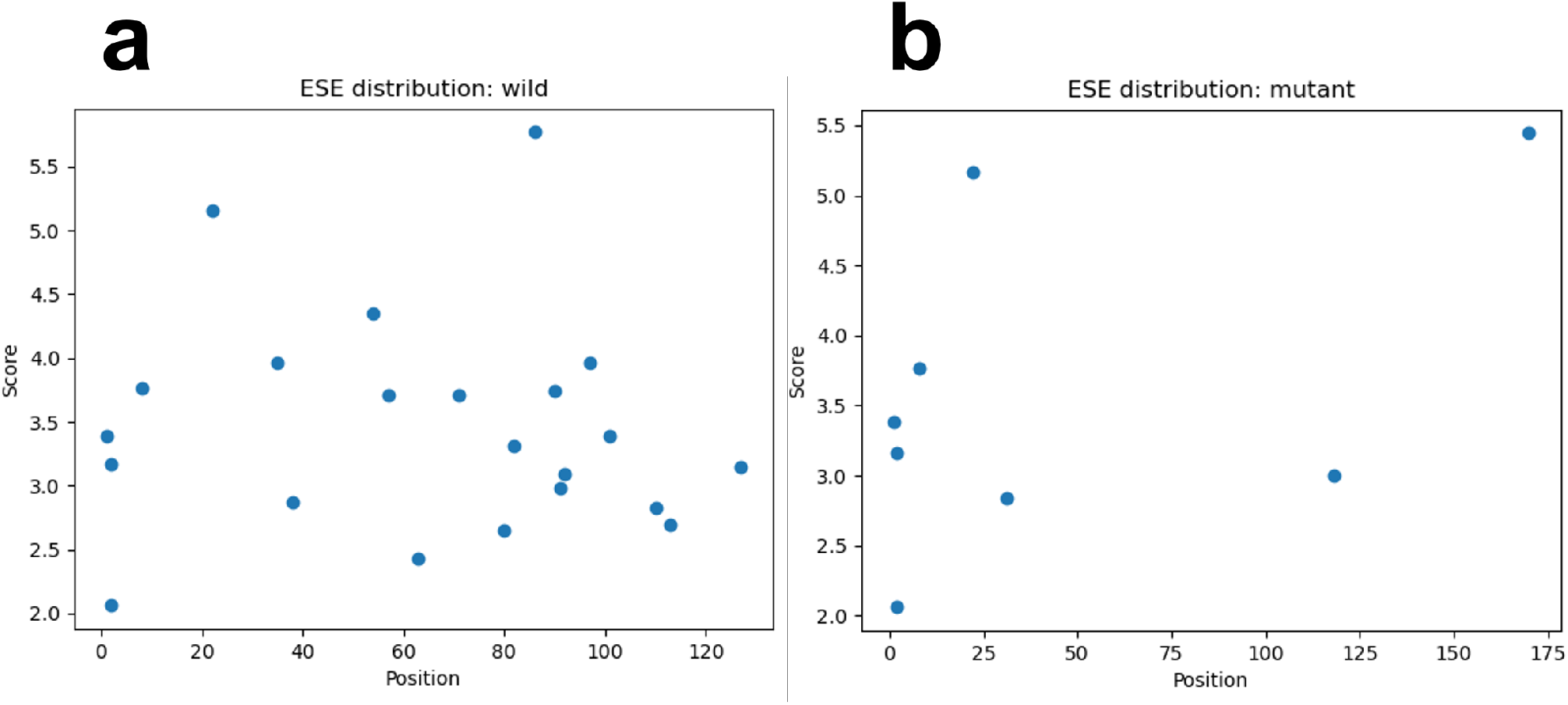
Predicted exonic splicing enhancers (ESEs) sites in the *scarlet* allele. (**a**) Predicted ESE sites in exon 5 of the wild-type *scarlet* allele. (**b**) Predicted ESE sites in exon 5 and the insertion region of the red-eye mutant *scarlet* allele.

**Figure S5.**
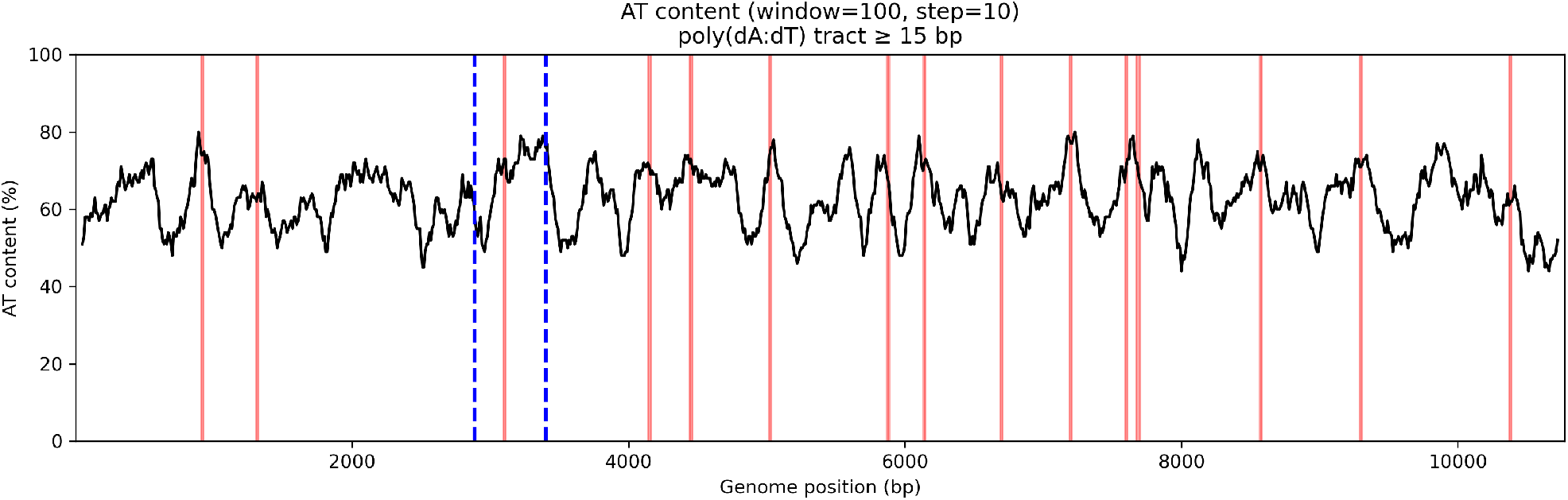
AT content across the *scarlet* gene. AT content was calculated using a 100-bp sliding window. Red lines indicate poly(dA:dT) tracts; blue dashed lines indicate the region deleted in the red-eye mutant.

## Supplementary Tables

**Table S1.**
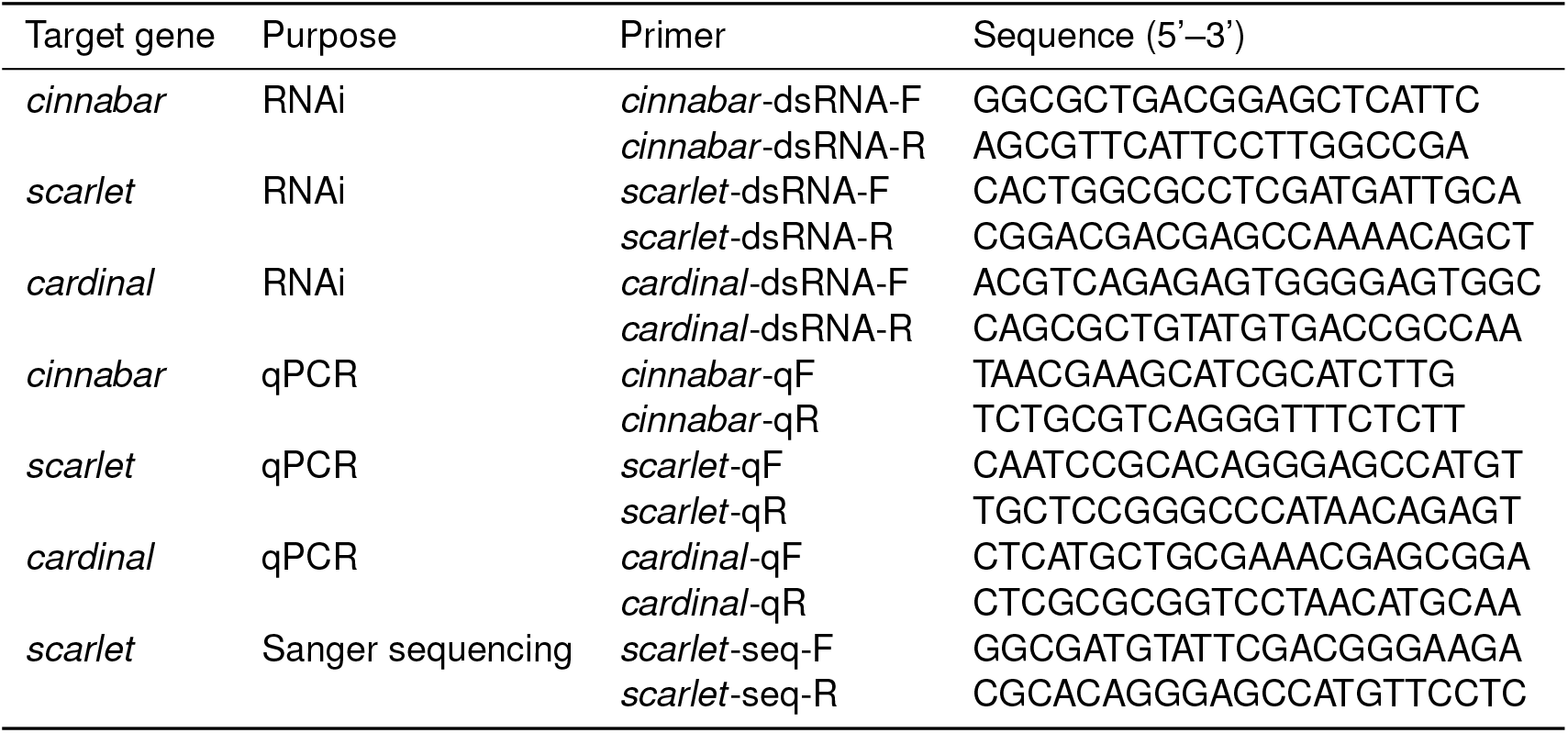
Primers used in this study.

